# In-channel cancellation: a model of early auditory processing

**DOI:** 10.1101/2022.11.27.518095

**Authors:** Cheveigné Alain de

## Abstract

A model of early auditory processing is proposed in which each peripheral channel is processed by a delay-and-subtract cancellation filter, tuned independently for each channel with a criterion of minimum power. For a channel dominated by a pure tone or a resolved partial of a complex tone, the optimal delay is its period. For a channel responding to harmonically-related partials, the optimal delay is their common fundamental period. Each peripheral channel is thus split into two subchannels, one that is cancellation-filtered and the other not. Perception can involve either or both, depending on the task. The model is illustrated by applying it to the masking asymmetry between pure tones and narrowband noise: a noise target masked by a tone is more easily detectable than a tone target masked by noise. The model is one of a wider class of models, monaural or binaural, that cancel irrelevant stimulus dimensions so as to attain invariance to competing sources. Similar to occlusion in the visual domain, cancellation yields sensory evidence that is incomplete, thus requiring Bayesian inference of an internal model of the world along the lines of Helmholtz’s doctrine of unconscious inference.

## I. Introduction

Acoustic information, transduced by the ear, is relayed through a series of processing stages within the auditory brain (brainstem, midbrain and cortex). The neural patterns carried by the auditory nerve reflect both the spectro-temporal features of the sound, and the fine temporal patterns of its waveform. Spectro-temporal features are resolved by cochlear filtering, coded by the slowly-varying firing rate across the array of auditory-nerve fibers, and projected to tonotopically-organized fields up to cortex. Fine temporal patterns are coded by the instantaneous firing probability within each fiber, and relayed by specialized neural circuits within the auditory brainstem up to the dendritic fields of the inferior colliculus.

A remarkable aspect of this circuitry is its high degree of fan-out, evident in cell numbers: ∼3000 inner hair cells, ∼100 000 neurons in cochlear nucleus (Hinojosa and Nelson, 2011), ∼100 000 000 in primary auditory cortex (Itoh et al., 2022). It is also evident in the diversity of cell types and patterns of response to sound at each stage. Some cells respond to onsets, others appear to smooth patterns over time or favor certain modulation rates. Others (the majority?) show patterns of response and selectivity that escape an easy explanation in terms of function. It has been argued that such a fan-out is useful in principle for metabolic and computational reasons (Barlow and Rosenblith, 1961; Gauthier, 2021). In any event, patterns transduced by the cochlea undergo many parallel transformations within the brain. This paper investigates a hypothetical neural circuit that might contribute to early auditory processing. The model posits, within each peripheral channel, a “cancellation filter” consisting of a neuron with an excitatory input and an inhibitory input, one of which is delayed relative to the other. The neuron performs a “gating” function: its properties are such that if a spike carried by the excitatory branch coincides with a spike on the inhibitory pathway, that spike is removed. Downstream processing has access to both the input of this filter and its output, as well as to the automatically determined delay parameter. The delay parameter is tuned automatically using information purely local to the channel (Fig. 1).

**Figure 1.**
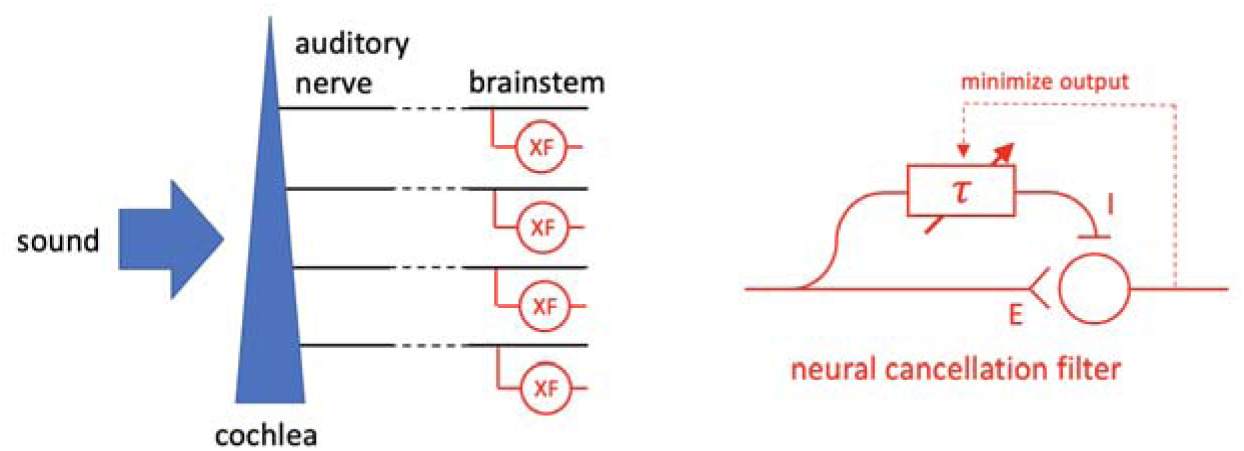
Left: each channel from the periphery is split into a direct pathway (black) and a “cancellation-filtered” pathway (XF, red). Subsequent processing can attend to either. Right: a cancellation filter consists of a gating neuron with a direct excitatory input and a delayed inhibitory input (or vice-versa). The delay is adjusted automatically to minimize the output.

Four arguments are developed in favor of this hypothesis. First, it can explain certain psychophysical results, such as the asymmetry of masking between a pure tone and a narrow band of noise (Hellman, 1972; Hall, 1997). Second, it may assist auditory scene analysis by calculating transforms of sensory input that are invariant to a certain type of interfering sound. Third, it extends an existing model (harmonic cancellation, de Cheveigné, 1993, 2021) to allow it to account for a wider range of phenomena. This extension follows the same logic by which the “modified equalization-cancellation model” (mEC) of Culling and Summerfield (1994) was derived from the earlier “equalization-cancellation” (EC) model of Durlach (1972). Fourth, it is plausible in terms of neural circuitry as, similar to the EC model, it involves fast excitatory-inhibitory interaction that is known to occur at several levels within the brainstem. These arguments are further developed in the Discussion.

A pure tone masker masks a narrowband noise probe less effectively than a narrowband noise masker masks a tone probe (Hellman, 1972, Hall, 1997). The difference in masking can reach 20 dB or more, a stark deviation from the standard “power spectrum model” of masking according to which a sound is detectable if it its power exceeds that of the masker within at least one peripheral frequency channel (Moore, 1995). Masking differences have also been observed between harmonic and noise-like maskers, as reviewed by de Cheveigné (2021): a harmonic masker is less effective than a noise-like masker of similar spectral envelope, suggesting an unmasking mechanism applicable to harmonic maskers.

In the spatial domain, an inter-aurally correlated masker (as produced by a spatially localized interfering source in an anechoic environment) is less effective than a masker that is inter-aurally uncorrelated. Some aspects of binaural unmasking are well accounted for by Durlach’s (1963) equalization-cancellation (EC) model that assumes that neural signals from both ears are differentially scaled and delayed (equalization), and then subtracted centrally (cancellation). Durlach’s (1963) EC model assumed, tacitly, that the same parameters (delay and scale factor) are applied to every peripheral channel. Refining that model, a more recent “modified EC” model allows for different equalization parameters in each channel (Culling and Summerfield, 1995; Breebart and Kohlrausch, 2001, Ackeroyd, 2004). With this modification, the mEC model can explain unmasking with maskers crafted to have different binaural properties (e.g. different interaural delay) in different frequency bands.

Similarly, the “harmonic cancellation” model of de Cheveigné (1993, 2021) assumes that a neural signal is delayed and subtracted from the non-delayed signal, with a delay equal to the period of an interfering sound, allowing correlates of a periodic (harmonic) masker to be suppressed. The original model assumed that the same delay was applied to every peripheral channel but, taking inspiration from the modified EC model, that assumption can be relaxed, resulting in the model described in this paper. With this modification, the model should account for unmasking observed with inharmonic stimuli obtained by stretching, shifting, or jittering the partials of a harmonic complex (Roberts and Brunstrom, 1998, 2001, Steinmetzger and Rosen, 2023). Inharmonic partials cannot be fit by a harmonic series across the spectrum, so there is no single delay that allows perfect cancellation, but they can be fit locally if their frequencies approximate a harmonic series over a restricted range.

Going one step further we can note that each individual partial is periodic and amounts to a one-component harmonic series with a fundamental equal to its period. The partial can be suppressed, within those channels that it dominates, by fitting a cancellation filter to that period. Thus, one expects some degree of unmasking for maskers that are spectrally sparse. The simplest example is a pure tone, as in Hellman’s study mentioned earlier.

The hypothesis of an automatically-tuned filter in each peripheral channel opens some interesting perspectives. First, it highlights the role of time-domain signal processing in the auditory brainstem, with the implication that auditory frequency selectivity is not entirely determined by cochlear frequency selectivity. The idea of a “second filter” dates back to Huggins and Licklider (1951), more recent incarnations being the lateral inhibitory network (LIN) of Shamma (1985) or the phase-opponency model of Carney et al. (2002).

Second, it emphasizes the role of invariance as a goal of auditory processing, in this case invariance to a certain class of background sounds (harmonic, quasi-harmonic or spectrally sparse). A downside of cancellation is that the integrity of the target is not guaranteed: certain of its features might be suppressed or distorted. Thus, cancellation needs to work hand-in-hand with a Bayesian or Helmholtzian process that can fit an incomplete or distorted target representation to an internal model (de Cheveigné, 2021).

Third, a by-product of the fitting process is an estimate of the period that dominates each channel. Whereas the original harmonic cancellation model involved a single period estimate that could serve as a cue to the pitch of a sound (de Cheveigné, 1998), the in-channel version offers multiple estimates. A stimulus for which period estimates differ across channels (e.g. an inharmonic complex) typically lacks a clear overall pitch. However, listeners may still be sensitive to pitch change (Demany and Ramos, 2005; Popham et al., 2018) that the present model might help explain.

The next section introduces the concepts needed to understand the model, and details of the simulations described in the Results section. Implications and limitations of the model are revisited in the Discussion section.

## II. Methods

### A. Model structure

The in-channel cancellation model requires both peripheral (cochlear) filtering, and central (neural) filtering. A simplified linear model of cochlear filtering is assumed here, and effects of non-linear transduction, compression, and stochastic neural coding are ignored for simplicity. The initial linear filter bank is followed by a delay-and-subtract cancellation filter that is a linear approximation of the excitatory-inhibitory operation schematized in Fig. 1 (right). The delay is estimated automatically and independently in each channel.

### B. Filters

Cochlear filtering is modeled using a gammatone filter bank (Holdsworth et al., 1988; Slaney, 1993) with bandwidths set to one equivalent rectangular bandwidth (ERB) according to estimates of Moore et al. (1983). Transfer functions of selected channels are plotted in Fig. 2, scaled so that their peak gain is one (0 dB). On a linear frequency scale, filters appear wider at high than low CF: bandwidth is roughly proportional to CF above 1 kHz, and roughly uniform over the lowest CFs. Each channel attenuates all but a narrow frequency region, so its output is relatively insensitive to the presence of signal features outside that region. However, at all frequencies the attenuation is finite.

**Fig. 2.**
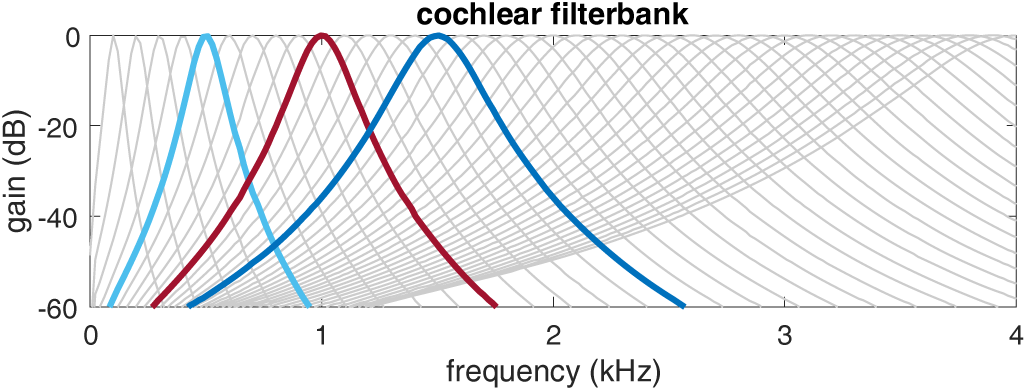
Transfer functions of selected channels of the gammatone filter bank. Channels with CF=0.5, 1, and 2 kHz are highlighted.

This selectivity can serve to “protect” a pure tone or narrowband target sound from a competing masker. This is illustrated in Fig. 3 (top) for a 1 kHz probe mixed with a wideband masker of equal RMS. Channels near 1 kHz are purely dominated by the probe, so the probe can be detected within these channels in spite of the masker. As another illustration (Fig. 3, middle), individual low-rank partials of a 200 Hz harmonic complex tone are isolated by channels with CFs near each partial’s frequency. Two ranges of CFs can be distinguished: below ∼1 kHz (5th harmonic), each partial appears to be perfectly isolated within a subset of channels close to its frequency, while intermediate channels respond to at most two neighboring partials. Above ∼1 kHz, individuals partials are less perfectly isolated (the peaks of the red curve do not reach one), and every channel seems to respond to more than two partials (the dotted line represents the power of the third-strongest partial).

**Fig. 3.**
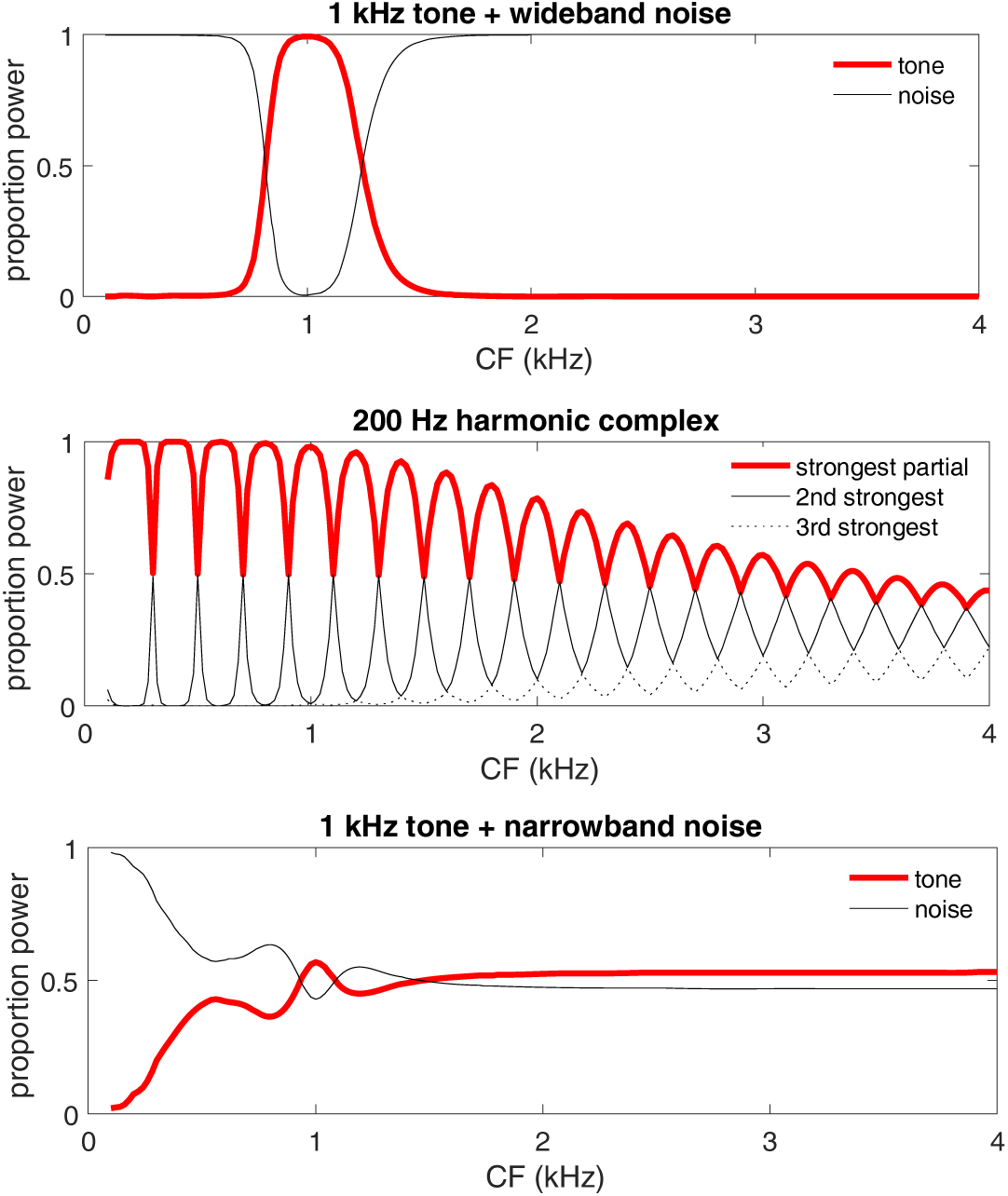
Proportion of power at the output of a cochlear filter bank as a function of CF for different parts of a stimulus. Top: mixture of a 1 kHz pure tone and wideband noise with equal RMS amplitude (over a 20 kHz bandwidth), red: proportion of power attributable to the pure tone, black: proportion attributable to the noise. Middle: 200 Hz harmonic complex tone. The red line represents the proportion of power attributable to the partial that dominates each channel. Continuous black: proportion attributable to the second strongest partial, dotted black: proportion attributable to the third-strongest partial. Bottom: mixture of a 1 kHz pure tone and a 0.5 ERB wide band of noise centered on 1 kHz, with equal RMS.

This parallels the classic distinction between “resolvable” and “unresolvable” partials of a complex (Moore and Gockel, 2011). That distinction is usually attributed to the presence or absence of salient ripples in the excitation pattern (e.g. Fig. 1 in Moore and Gockel, 2001), but Fig. 3 (middle) offers a different perspective: a partial is resolvable if it dominates at least one peripheral channel (i.e. within that channel the power of all other partials is below some threshold). This might, for example, be a condition for the estimation of that partial’s frequency based on time-domain neural patterns (Srulovicz and Goldstein, 1983).

Figure 3 top and middle illustrate situations where peripheral filtering can protect a probe sound from a competing background. In contrast, Fig. 3 (bottom) shows a situation where a peripheral filter is of little help. The stimulus here is a mixture of a pure tone at 1 kHz and a narrowband noise centered at 1 kHz with 0.5 ERB bandwidth (∼60 Hz) and equal RMS amplitude. No channel is clearly dominated by the pure tone. This leads us to expect a high threshold for detecting a pure tone probe in a narrowband noise, as is indeed observed in psychophysical experiments.

The second component of the model is the cancellation filter, modeled as a delay-and-subtract filter with impulse response (Fig. 4 left), approximation of the “neural filter” of Fig. 1 (right). The transfer function has zeros at all multiples of (Fig. 4 right), implying that attenuation is infinite at those frequencies.

**Fig. 4.**
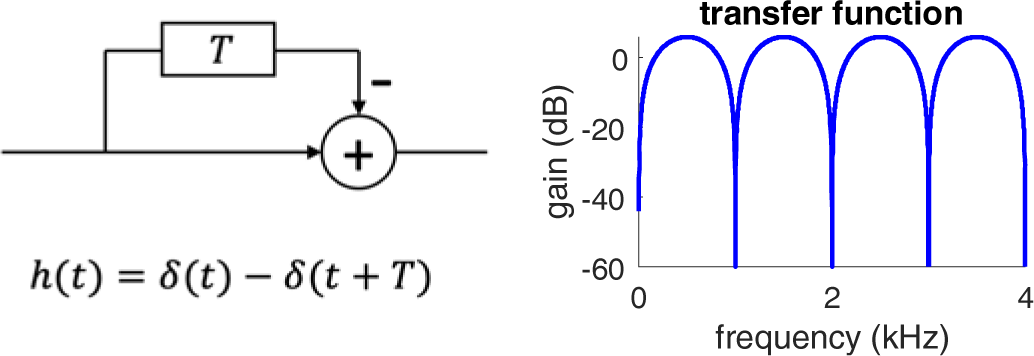
Left: schema and impulse response of a cancellation filter. Right: transfer function for =1 ms.

If the cancellation filter is applied to the output of a gammatone filter channel, the transfer function of the cascade is the product of their transfer functions, as illustrated in Fig. 5 for three channels with CFs in the vicinity of 1 kHz. The compound filter inherits spectral filtering properties of its two parts: in addition to reduced gain at frequencies remote from its CF, it has infinite attenuation at 1 kHz. The output of this compound filter is invariant to the presence, or absence, of spectral power at precisely 1 kHz.

**Fig. 5.**
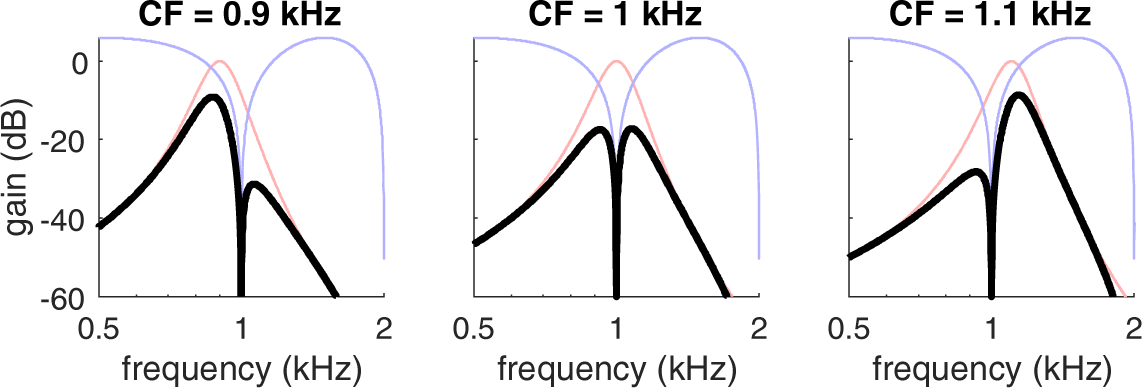
Transfer function of the cascade of a gammatone filter and a cancellation filter with a delay parameter =1 ms for three CFs (as indicated). The thin colored lines are the transfer functions of the component filters (red: gammatone, blue: cancellation).

### C. Finding

The in-channel cancellation filter requires deciding the delay in each channel . Here, we make the important assumption that is determined within each channel on the basis of the signal within that channel. This is achieved as schematized in Fig. 6, by searching for the non-zero value of that minimizes the power at the output of the cancellation filter relative to its input. This is analogous to well-known techniques for period estimation such as AMDF (average magnitude difference function, Ross, 1976) or YIN (de Cheveigné and Kawahara, 2002). For a periodic sound, the minimum is zero, in the presence of a probe it is non-zero, however even in that case the delay estimate is accurate as long as the probe is weak (which should be the case at detection threshold).

**Fig. 6.**
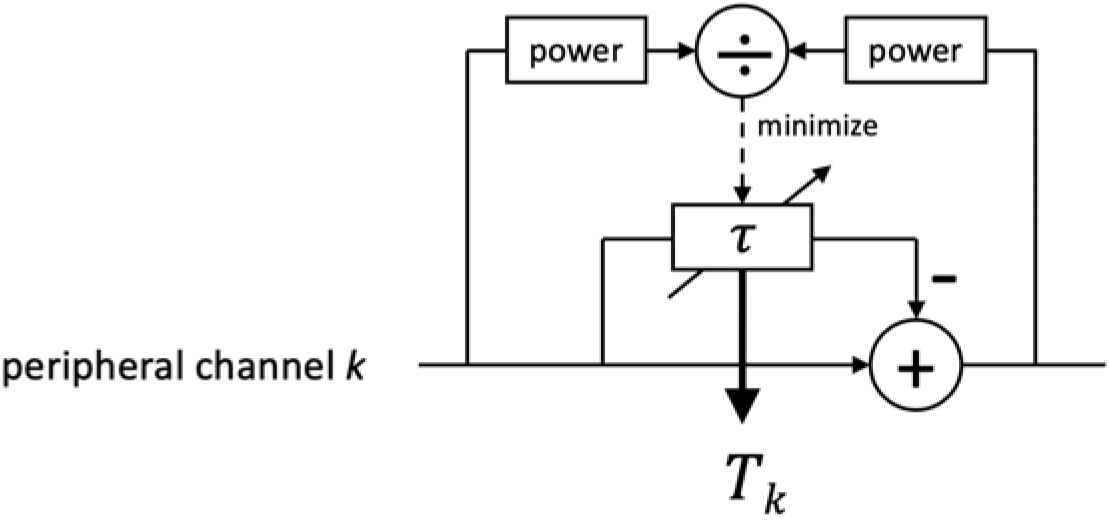
Automatic determination of the delay within each channel . The delay parameter is varied in search for the smallest ratio of cancellation filter output power to input power.

A couple of details need elaborating. First, input and output power must be smoothed to get a stable estimate of their ratio, and the smoothing window must be long enough to suppress irrelevant fluctuations, yet short enough to track period changes in a non-stationary signal. In the simulations the window size was set to 30 ms. Second, for a purely periodic signal the output/input ratio is zero for the period and all its multiples, implying an infinite number of equally valid candidates for . For definiteness, it may be convenient to choose among candidates the first for which the power ratio is below some threshold (see de Cheveigné and Kawahara, 2002 for a rationale and more details). This parameter may affect but it has little impact on the ability of the filter to suppress the periodic background.

Figure 7 plots the inverse of the automatically-determined delay T_k_ within each channel as a function of CF for three stimuli: a 1 kHz pure tone (left), wideband noise (center), and narrowband noise (right). The parameter 0 was set to 0 in each case. For the pure tone, the estimate is 1/T_k_ = 1 kHz within all channels. For wideband noise, it is close to CF for every channel, suggesting that it reflects the noise-excited ringing pattern within each channel. For the narrowband noise, the inverse estimate follows CF at low CFs and stabilizes near 1 kHz at high CFs. These examples illustrate how the delay T_k_ is automatically chosen for three common stimuli.

**Fig. 7.**
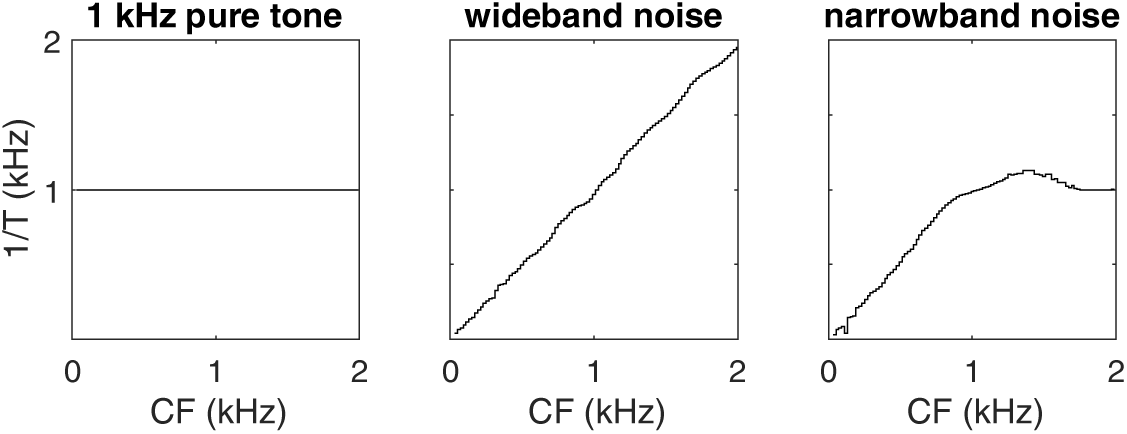
Inverse of estimated delay T for each channel as a function of its CF for three stimuli. For the pure tone (left), all channels yield 1/T = 1 kHz. For wideband noise (center), the value is approximately 1/T =CF. For narrowband noise (right) it follows 1/T =CF at low frequencies and is close to 1/T = 1 kHz at high CFs.

### D. Simulations

Peripheral filtering was simulated with a gammatone filter bank as explained earlier. Either one channel (CF = 1 kHz) or multiple channels spaced 20 Hz from 30 Hz to 2000 Hz (or 4000 Hz, as indicated by the axis label) were simulated. The cancellation filter was implemented as a simple finite impulse response (FIR) filter, with fractional delays handled using linear interpolation. Spectro-temporal excitation patterns were calculated by smoothing the instantaneous power within each peripheral channel (before or after cancellation) with a 30 ms boxcar window. The color code of time-frequency plots (“spectro-temporal excitation patterns”) represents values in dB relative to their maximum. Period estimation was performed using the YIN algorithm that implements the algorithm sketched in Fig 6 (de Cheveigné and Kawahara, 2002, http://audition.ens.fr/adc/sw/yin.zip), with a threshold 0 equal to 0.

Stimuli consisted of a probe (pure tone or narrowband noise pulse) superimposed on a masker (pure tone, sweep frequency pure tone, harmonic complex, inharmonic complex, wideband noise or narrowband noise). A pure tone probe was produced by applying a 30 ms window with 10 ms raised-cosine onset and offset ramps to a 1 kHz sinusoid. A narrow-band noise probe was produced by applying the same temporal window to narrowband noise obtained by filtering wideband Gaussian noise with a 4-th order gammatone filter with center frequency 1kHz and bandwidth 0.5 ERB (∼66 Hz). The onset of a probe was always temporally centered on the masker.

The pure tone masker was obtained by applying a 0.5 s window with 10 ms onset and offset ramps to a sinusoid of frequency 1 kHz. The sweep tone masker was similar but its frequency was swept on a logarithmic scale from 0.5 kHz to 2 kHz over its duration (0.5 s). The harmonic complex masker was produced by adding pure tones with frequencies multiple of 200 Hz up to 5kHz in cosine phase, with the same temporal window. Inharmonic complex maskers were obtained by either shifting all these frequencies downwards by 50 Hz (shifted condition), or jittering these frequencies with values drawn uniformly between - 100 Hz and +100 Hz, the draw being renewed until no partials were closer than 30 Hz (jittered condition). The wideband noise masker was Gaussian noise, the narrowband noise masker was obtained by filtering the same with a gammatone filter centered on 1 kHz with bandwidth 0.5 ERB (∼66 Hz). Both were temporally windowed as the other maskers. Stimuli are scaled between [-1, 1], with units that bear no relation (other than proportionality) to sound pressure, basilar membrane motion, or instantaneous firing probability. The nominal level of a masker or probe is its RMS level before windowing.

## III. Results

Three examples are considered: the masking level difference between pure tone and narrowband noise (Hellman, 1972), masking of a narrowband probe by a sweep-frequency masker (Smoorenburg and Coninx, 1980), and masking by harmonic and inharmonic complex tones (e.g. Roberts and Brunstrom, 2001).

### A. Example 1: masking level difference between pure tone and narrowband noise

This example is chosen for its simplicity and relevance to the asymmetry of masking between pure tone and narrow-band noise observed by Hellman (1972; Hall, 1997; Schroeder et al., 1979). For a pure tone masker, according to Fig. 4 (right) the output of the cancellation filter should be zero, assuming the delay T_k_ is accurately chosen. Actually, this is not quite the case because the stimulus is of finite length: there is a transient at both onset and offset, but the part intermediate between them is zero (Fig. 8 top right). The situation is different for a narrowband noise masker (Fig. 8 bottom right), for which the compound filter fails to cancel any portion. In both examples, the delay T_k_ was chosen automatically as described earlier. For the pure tone masker this delay equals 1 ms for all peripheral channels, for the narrowband masker it varies as in Fig. 7 (right). The conclusion to be drawn from this simulation is that these two maskers differ in that the cancellation filter can effectively suppress one but not the other. This is corroborated by Fig. 9 that shows the spectro-temporal excitation patterns for these same stimuli before and after cancellation. The response of all channels is suppressed for most of the duration of the stimulus for the pure tone (top right), but not the narrowband noise (bottom right).

**Fig. 8.**
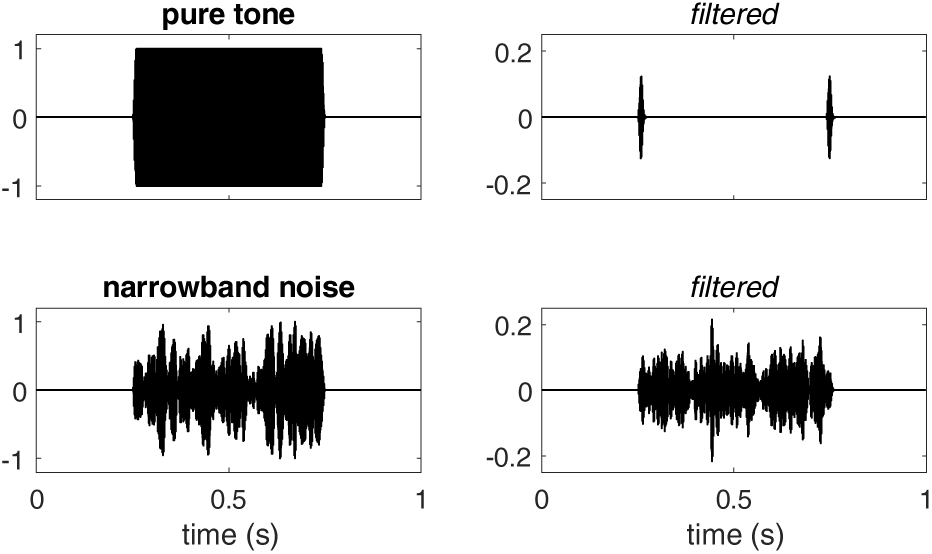
Effect of the in-channel cancellation filter within the CF=1 kHz channel for two stimuli. Top: 1 kHz pure tone, bottom: narrowband noise centered on 1 kHz (width 0.5 ERB). Left: stimulus, right: compound filter output. Stimuli are 500 ms in duration and shaped with 10 ms raised-cosine onset and offset ramps.

**Fig. 9.**
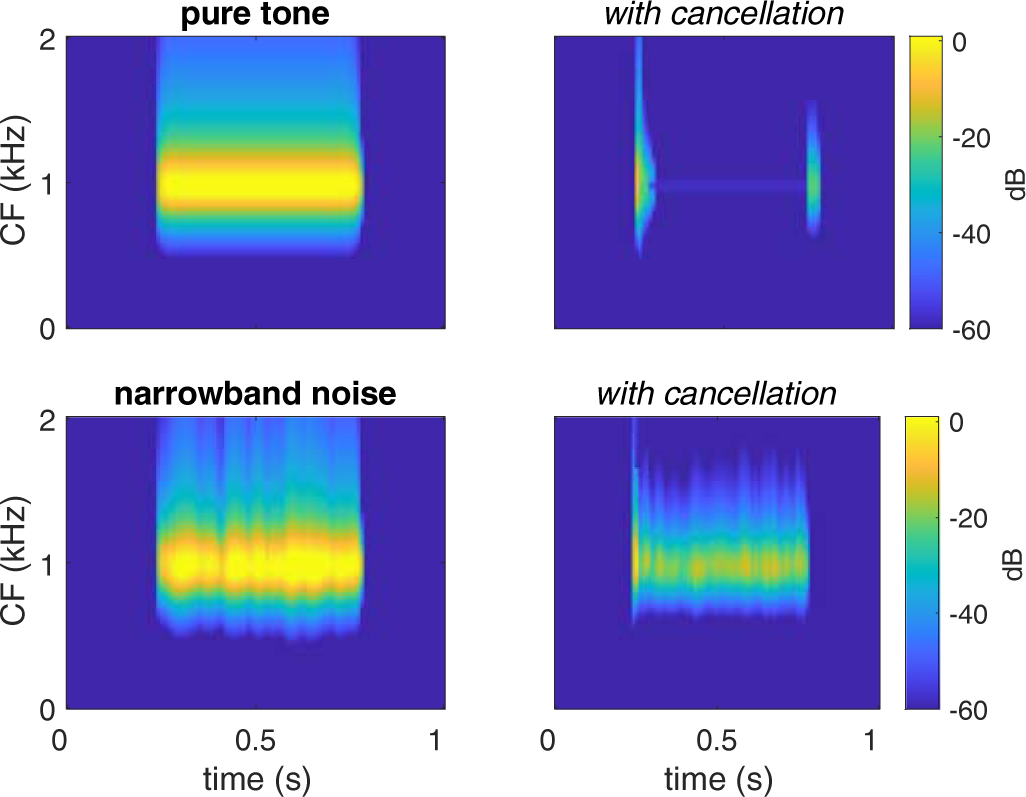
Spectro-temporal excitation patterns for the same stimuli as in Fig. 8, before (left) and after (right) cancellation. The auditory system is assumed to have access to spectro-temporal excitation patterns both with and without cancellation.

Figure 10 is similar to Fig. 9, with the addition of a 30 ms narrowband probe temporally centered on the pure tone (top) or narrowband noise (bottom) masker. The amplitude of the probe was 10 dB below that of the masker, which according to Hellman (1972) should make it suprathreshold for a pure tone masker and subthreshold for a narrowband noise masker. Its presence is not visually detectable in the absence of cancellation (left). With cancellation, it is prominent for the pure tone masker (top right) but not the narrowband noise masker (bottom right).

**Fig. 10.**
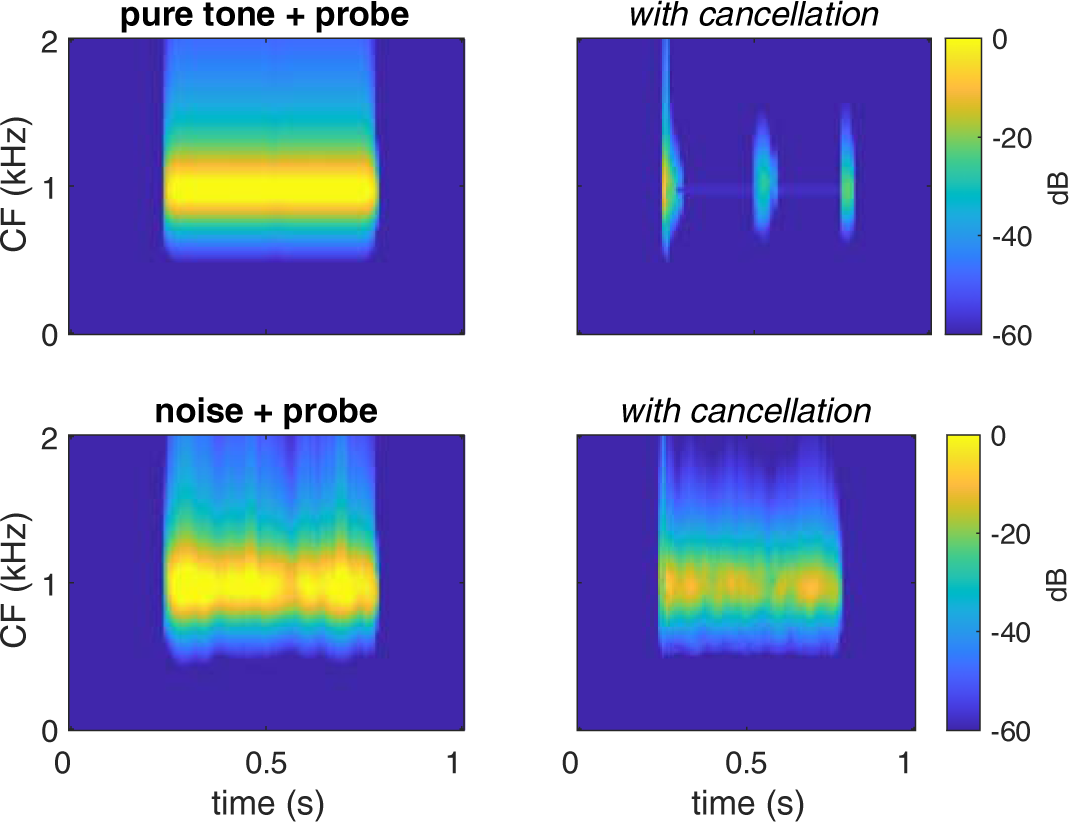
Spectro-temporal excitation patterns before (left) and after (right) cancellation for a probe added to a masker. The masker was a 1 kHz tone (top) or a narrowband noise of width 0.5 ERB (bottom). The probe was a 30 ms pulse of narrowband noise temporally centered within the masker, with an amplitude -10 dB relative to the masker.

This is qualitatively consistent with the observation, made by Hellman (1972) and others, that a pure tone masker is much less potent than a narrowband masker of similar amplitude. Here, this emerges as a consequence of the availability within each peripheral channel of a self-tuned cancellation filter (Fig. 1). However, other explanations have been put forward that are reviewed in the Discussion (Future Directions).

### B. Example 2: frequency sweep tone masker

Smoorenburg and Coninx (1980) found that a pure tone probe was more effectively masked by a sinusoidal masker with a logarithmically modulated frequency (sweep) than by a pure tone, for a probe frequency matching the frequency of the pure tone or the instantaneous frequency of the sweep at the probe time. The masking level difference was up to 20 dB. The standard power-spectrum model of masking (Moore 1995) would predict less masking for the sweep because it contributes only briefly to power within the channel occupied by the probe. Edwards and Viemeister (1997) found similar effects with other forms of frequency modulation.

The previous simulation was repeated, replacing the narrowband noise masker with a sinusoidal masker with frequency swept from 500 Hz to 2 kHz in 0.5 s (sweep rate 4 octaves/s). A 1 kHz pure tone probe was added, temporally centered on the pure tone or sweep tone masker. Figure 11 shows the spectro-temporal excitation pattern after cancellation filtering. For the pure tone masker (Fig. 10 left), the presence of the probe is visually salient (as in Fig. 9 top right). For the sweep tone masker, cancellation is imperfect due to the non-stationarity of the frequency-modulated masker, so the probe is barely perceptible visually after cancellation (Fig. 10 right). This is in qualitative agreement with the observation made by Smoorenburg and Coninx (1980) that a sweep tone is a stronger masker than a pure tone. Again, other explanations have been put forward that are reviewed in the Discussion.

**Fig. 11.**
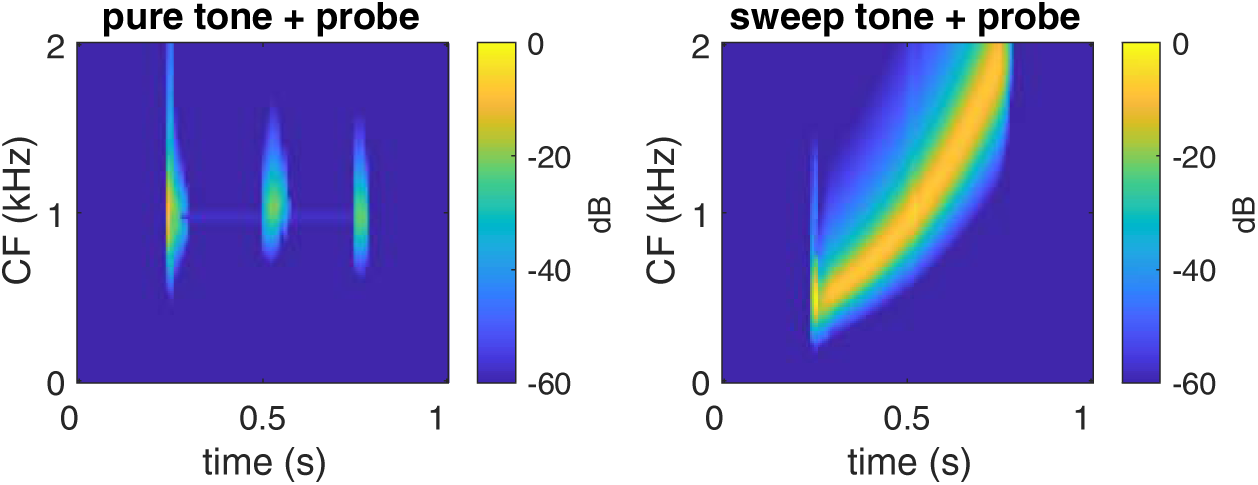
Spectro-temporal excitation patterns before (left) and after (right) cancellation for a short pure tone probe added to a frequency sweep masker. Cancellation allows the probe to emerge (visually) from the pure tone but not the sweep.

### C. Example 3: harmonic and inharmonic maskers

A similar masking level difference has been found between harmonic complex maskers and inharmonic or noise-like maskers (Treurniet and Boucher, 2001; Gockel et al., 2003; Deroche and Culling, 2011): masker harmonicity (or periodicity in the time domain) is associated with low detection thresholds. An advantage is also found for suprathreshold perception, for example of vowel timbre (Summerfield and Culling, 1992; de Cheveigné et al., 1997, see de Cheveigné, 2021 for a review). Surprisingly, however, a similar advantage has also been reported for maskers that are not perfectly harmonic, either because the partials of a harmonic complex were all shifted by the same frequency, or because they were jittered by random amounts, in both cases making the complex inharmonic (Roberts and Brunstrom, 1998, 2001; Deroche et al., 2014; Steinmetzger and Rosen, 2023).

Figure 12 (top) shows spectro-temporal excitation patterns before (left) and after (right) cancellation, for a 200 Hz harmonic complex tone, and an inharmonic complex obtained by applying random jitter to the frequencies of the harmonic complex (see Methods). The jittered complex is attenuated in certain spectral regions (their position depends on the values drawn for the jitter), albeit not as much as the harmonic complex. Figure 12 (bottom) shows the amount of attenuation as a function of CF for Gaussian noise (black), a harmonic complex (blue), a harmonic complex with partial frequencies shifted by -50 Hz (green) and a jittered harmonic complex. Considerable attenuation is attained for both inharmonic maskers, relative to a noise-like masker, which might explain why inharmonic maskers share some of the properties of harmonic maskers (Roberts and Brunstrom, 1998, 2001; Steinmetzger and Rosen, 2023).

**Fig. 12.**
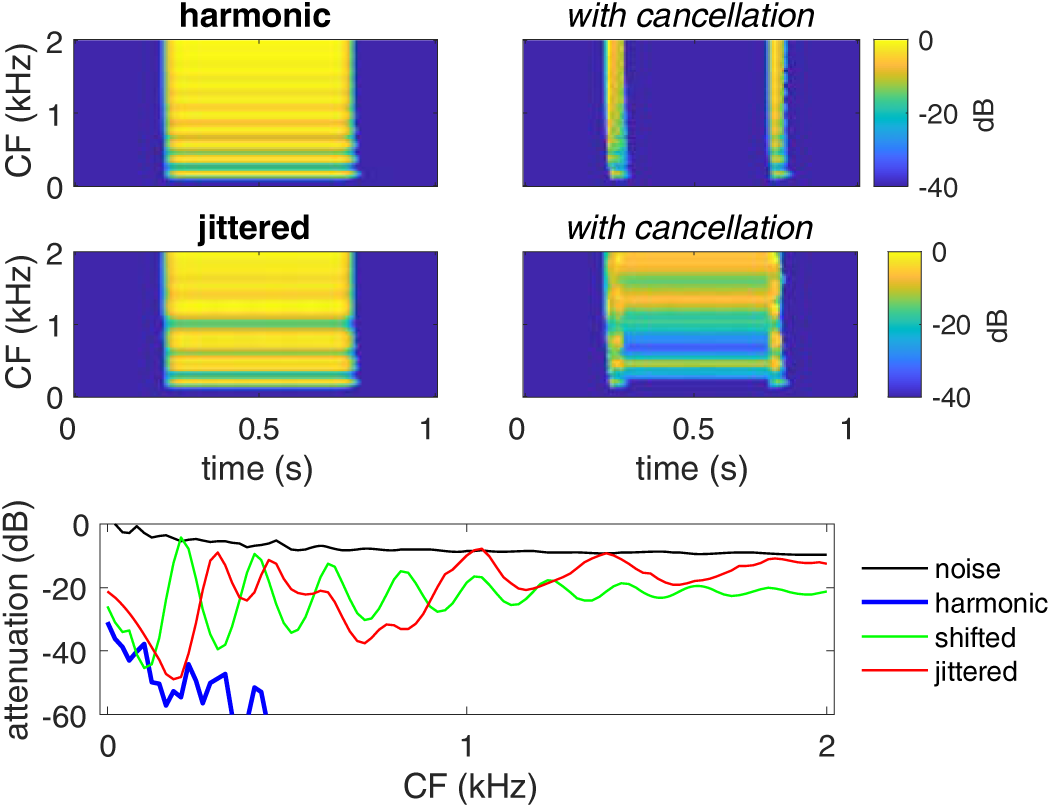
Top: spectro-temporal excitation patterns before (left) and after (right) cancellation for harmonic and and inharmonic maskers (no probe). Bottom: ratio of output/input power averaged over the central portion of the stimulus (0.3 to 0.7 s) for noise, harmonic, shifted (−25% of f0) and jittered complexes. Shifted and jittered complex maskers can be cancelled better than noise, but less well than a harmonic masker (blue, most values below -60 dB).

Note that the values in both plots depend on the search range for the best delay (or equivalently the range of delays applicable in the filter) in each channel, here arbitrarily limited to 0– 16.7 ms (inverse of 60 Hz). If the range had been 0–20 ms (inverse of 50 Hz) or wider, the search would have discovered the true period of the shifted complex, resulting in the same attenuation as for the harmonic masker.

## IV. Discussion

In-channel cancellation is a hypothetical neural process by which each peripheral auditory channel is split into two branches, one filtered by an automatically-tuned cancellation filter, the other left unfiltered.

Within the filtered branch, a pure tone masker is perfectly cancelled, thanks to its periodicity, and so adding a probe, even faint, produces a non-zero output that is salient on the otherwise zero background of the cancellation filter output (Fig 10 top right, Fig 11 left). A narrowband noise or sweep tone masker are not so well cancelled, and thus the addition of a probe is not noticed in that case, consistent with the results of Hellman (1972) and Smoorenburg and Coninx (1980), and similar reasoning can be applied to explain masking differences between harmonic and non-harmonic maskers. Other explanations have been put forward for those effects that are discussed in subsection Future Directions.

A major goal of auditory processing is to ensure invariance, both to irrelevant sound dimensions for the purpose of classification, and to competing sound sources for the purpose of auditory scene analysis. Cochlear frequency resolution (Moore, 1995) and temporal resolution (Plack et al 1990) can be interpreted in this light, as they allow interference-dominated spectral or temporal intervals to be ignored, the remainder being invariant to the interference. For example, cochlear filtering isolates a narrowband target from the presence of off-band energy (Fig. 3 top), and in-channel cancellation extends this to an on-frequency tonal masker for which cochlear filtering does not suffice (Fig. 3 bottom). Cochlear filtering and cancellation both contribute to invariance by suppressing the masker, but target features that fall within discarded channels or on zeros of the cancellation filter (Fig. 4, right) are distorted or lost. One must therefore assume that cancellation is followed by a process of unconscience inference (Helmholtz, 1867) by which a perceptual object can be inferred from the available (albeit incomplete) information thanks to an internal model (de Cheveigné, 2021).

From a different perspective, in-channel cancellation is related to predictive coding (Barlow and Rosenblith, 1961; Friston, 2018), as the sensory pattern within each peripheral channel at each instant is predicted from its value r in the past, and the two subtracted. The outcome of this coding is a delay parameter T_k_ that contributes to characterize the stimulus, and an “error signal” from which a weaker concurrent target can be inferred. This is also similar in principle to linear predictive coding (LPC) (Atal, 2006).

The availability of a mechanism that can reduce the masking power of certain sounds implies for those sounds a property of transparency, that may be beneficial in a musical context that involves building a complex sound structure with multiple elements (polyphony). In the context of perceptual coding of audio signals, that property (referred to in this context as “tonality”) must be taken into account as it enhances the detectability of quantization noise (Schroeder et al, 1979; Johnston, 1988; Estreder et al., 2023).

The in-channel cancellation model is an extension of the earlier harmonic cancellation model of de Cheveigné (1993, 2021). The new model is more flexible than the old, and can account for a wider range of phenomena. It might seem more complex and less parsimonious because it involves more parameters (the value of T_k_ in each channel).

However, the T_k_ are chosen automatically and thus do not constitute “free” parameters (they cannot be adjusted to improve the fit to a data set). One might argue instead that the in-channel model is simpler because it is implemented locally within each channel, with no need for a cross-channel mechanism to derive a common T across channels and distribute that estimate to each channel. The relationship is analogous to that between the original and modified EC models of binaural unmasking (Durlach, 1963; Culling and Summerfield, 1994; Breebart and Kohlrausch, 2001; Ackeroyd, 2004).

In the neural circuit of Fig. 1 (right), the spike train carried by the inhibitory branch controls a “thinning” process applied to the spike train carried by the excitatory branch. This is similar to the excitatory-inhibitory interaction thought to occur in the Lateral Superior Olive (LSO) (Beiderbeck, 2018; Franken et al 2021) for the purpose of binaural interaction, in a process similar to the EC model. There, a spike train from the contralateral ear controls (after a delay to compensate for interaural delay related to the spatial characteristics of an interfering source) a thinning process applied to a spike train from the ipsilateral ear. The similarity in principle (inhibitory thinning process) and goal (interference cancellation) is an argument in favour of the present model. It is remarkable that removing a subset of spikes from a spike train can have an effect similar to the subtraction operation in the cancellation filter (Fig. 4, left) or EC model. Previous modelling has shown its effectiveness for recorded auditory-nerve patterns (Palmer, 1990) or simulated spike trains (de Cheveigné, 1993, Guest and Oxenham, 2019). In addition to LSO, there are multiple sites within the auditory brainstem where such excitatory-inhibitory interactions might in principle occur, from dendritic fields of the cochlear nucleus to dendritic fields of the inferior colliculus (see de Cheveigné, 2021 for a review).

### A. Future directions

The in-channel cancellation hypothesis raises a number of questions that are worth pursuing, either to challenge the hypothesis, or to explore avenues that it suggests.

#### 1. Does it work for speech?

Considerable efforts have been invested in assessing whether segregation mechanisms revealed using synthetic stimuli (“double vowels”), are also effective for “real-world” stimuli such as speech (e.g. Deroche et al., 2017; Steinmetzger and Rosen, 2023). A priori, there are reasons to expect that harmonic cancellation might not be effective, as the non-stationarity of the speech signal is likely to hinder both the reliability of masker period estimation, and the effectiveness of cancellation.

The situation might be slightly better with in-channel cancellation, as it offers flexibility to exploit “islands” of harmonicity within the spectro-temporal map, but this needs experimental confirmation. Figure 13 (left) shows the spectro-temporal excitation pattern in response to a short speech phrase (“Wow Cool!”). Fig. 13 (right) shows the degree of suppression afforded by in-channel cancellation (yellow: no suppression, blue: strong suppression). Clear suppression is evident in a small number of time-frequency pixels (right), which fortunately correspond to pixels of high amplitude in the spectro-temporal excitation pattern (left). Within certain pixels the attenuation reaches 20 dB or more, which might make it easier to hear features of a weak target (e.g. other voice) that happen to fall within those pixels. Whether this can translate into a benefit in terms of intelligibility in a “cocktail party” situation remains to be determined.

**Fig. 13.**
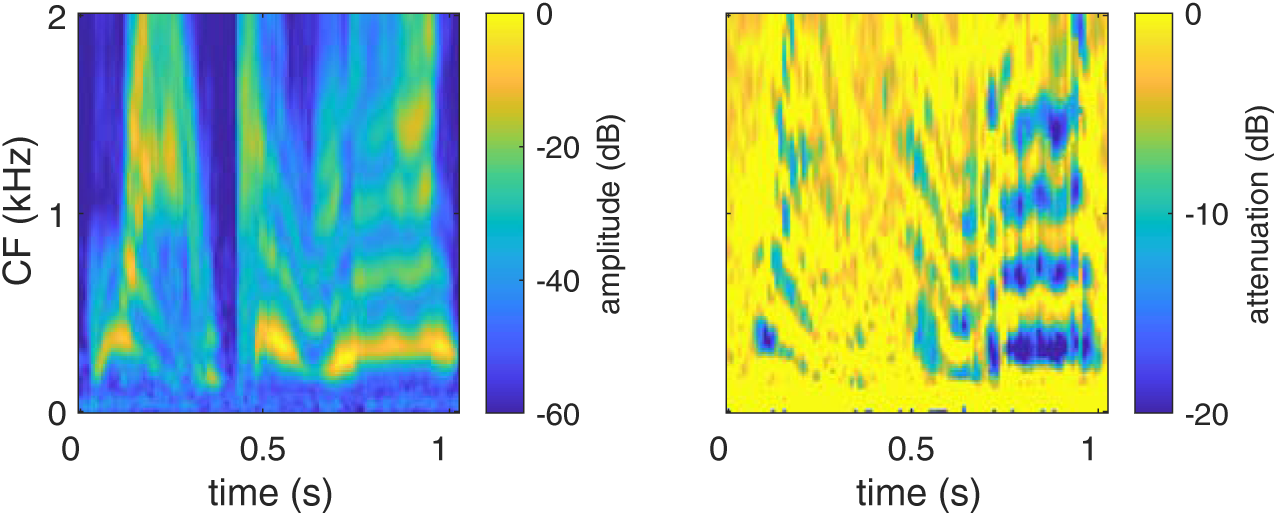
Left: spectro-temporal excitation pattern in response to a short sentence (“Wow, Cool!”). Right: attenuation provided by the in-channel cancellation filter (ratio of output power to input power for each channel and time step).

#### 2. In-channel cancellation, envelope fluctuations, and the modulation spectrum

Other explanations have been proposed for the masking phenomena described earlier. An obvious difference between a pure tone (or harmonic) masker and a narrowband noise (or non-harmonic) masker, is that the former has a relatively smooth temporal envelope while the latter fluctuates. Fluctuations evoked by the masker might confuse detection of the probe. For example, Edwards and Viemeister (1997) pointed out that the sweep tone in Smoorenburg and Coninx (1980) produces a momentary increase in power within the channel centered on the probe, temporally coincident with the increase expected from the probe. Verhey (2002) likewise explained the reduced masking by a narrow bandpass noise masker of a wider bandpass noise probe (of which the narrowband-on-pure tone condition of Hellman 1972 is a limit case) on the basis of a modulation filterbank (Dau et al., 1991). Modulation cues (e.g. roughness or beats) have also been invoked to explain f0-based segregation of concurrent vowels (e.g. Culling and Darwin, 1994), or asymmetric masking between harmonic and inharmonic sounds (e.g. Treurniet and Boucher, 2001, although see de Cheveigné et al., 1997, Deroche et al., 2014).

More work is required to decide between these models, or possibly unify them. A cancellation filter is a sensitive mechanism to detect envelope fluctuations of a sinusoidal or harmonic carrier, and might conceivably contribute to the implementation of an internal modulation filter bank (Viemeister, 1979; Dau et al., 1991; Jepsen et al., 2008). Indeed, the neural cancellation filter has the same structure as the same-frequency inhibitory-excitatory (SFIE) circuit that Nelson and Carney (2004) proposed to explain amplitude-modulation tuning in the inferior colliculus.

#### 3. Pitch and pitch change

Cancellation is closely related to autocorrelation, and its optimal delay parameter T has been proposed as a cue to pitch (de Cheveigné, 1998). The in-channel model is particular in that it offers multiple delay estimates T_k_, each corresponding to the period of a partial, or of a group of harmonically-related partials. Their perceptual relevance, if any, is worth investigating. One possibility is that they serve as cues to a pitch local to a spectral region (accessible to consciousness via attention to that spectral region). Another is that they underlie “partial pitches” with perceptual reality at a sub-attentive level (analogous to the “spectral pitches” invoked by Terhardt, 1974). A third is that they might be the basis for frequency shift detectors, by which one might perceive upward or downward shift of a set of partials as a change in pitch, even if those partials are not harmonically related, and do not evoke a clear sensation of pitch in the absence of the shift (Demany and Ramos, 2005, Demany et al., 2011; Chambers and Pressnitzer, 2014; McPherson et al., 2018; Demany and Semal, 2018).

A possible approach to pitch shift detection is schematized in Fig. 14 for one channel. The period T_k_ within a channel is manifest as the position of a minimum in output across an array of cancellation filters. The direction of frequency change can then be found by monitoring the array to the immediate left of that minimum (shorter delays) and noting whether the filter output decreased or increased. A decrease indicates an upward frequency shift, an increase indicates a downward shift. A reason to consider this cue is that filter output power is a prothetic quantity that might possibly be easier to remember than the metathetic quantity position of the minimum, or possibly easier to aggregate across channels, either by an averaging or voting mechanism (e.g. to explain McPherson et al., 2018, or Allik et al., 1989), or by selective attention to channels for which there is change (e.g. to explain Demany and Ramos, 2005). This hypothesis remains to be investigated.

**Fig. 14.**
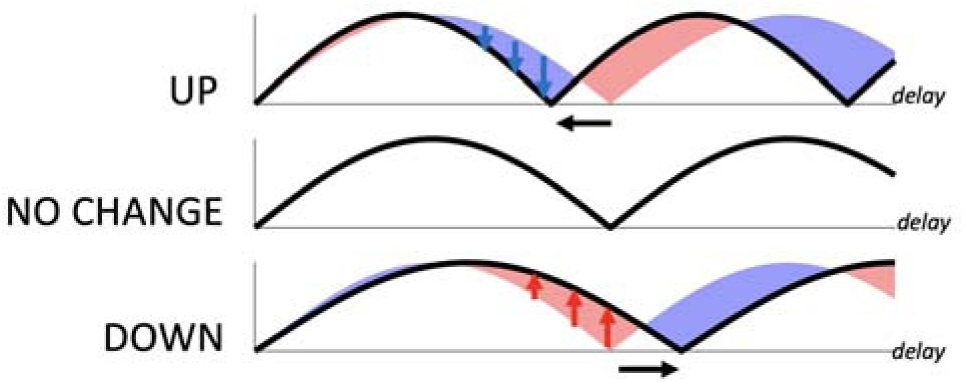
Schematic principle of an in-channel frequency shift detector. Each plot represents the amplitude at the output of a cancellation filter as a function of its delay parameter (or across an array of such filters indexed by delay) at two moments in time. For an upward change in frequency (top), the amplitude drops to the left of the original minimum (shorter delay), for a downward change (longer delay) it increases. This detector is instantiated independently in all channels and shift cues aggregated across channels.

#### 4. Spectro-temporal fan-out in the brainstem

The Introduction pointed out the remarkable fan-out of the auditory system. Multiple transforms of sensory input are useful to discover boundaries that ensure invariance to irrelevant dimensions (Duda et al., 2012; Gauthier et al., 2021). Random transforms suffice, as long as they are sufficiently numerous and include convolution and non-linearity, but there is a benefit to including a priori useful transforms within the mix. In-channel cancellation is a priori useful in that it ensures invariance to a class of potential interfering stimuli (harmonic and/or spectrally sparse). It is a “good thing to have” for the auditory brainstem.

The cancellation filter (Fig. 4) is an FIR filter, one of the simplest, and it is worth considering whether a wider class of signal processing operations might profit from the synergy between cochlear and neural filters. A possible direction is sketched here. Using a sampled-signal notation for convenience, the cochlear filter bank can be approximated as a M-column matrix M of finite impulse responses of order N. Filtering amounts to multiplying the N-column matrix X of delayed stimulus signals (delays 1· N) by the matrix M to obtain the matrix Y= XM of cochlear-filtered signals (one channel per column). If the matrix M allows an inverse, (i.e. if its rank is larger than N), multiplying Y by this inverse would reconstruct X= YM^-1^, i.e. the acoustic waveform (a single column of X) could be reconstructed as a weighted sum of cochlear filter outputs. Furthermore, supposing that a task can be addressed by applying some filter of impulse response h to the acoustic waveform, the output of that filter is available by multiplying Y by M^-1^h. In other words, any filter (of order at most N) applicable to the acoustic stimulus can be implemented within the auditory brain. The filtered signal is merely a weighted sum of cochlear filter outputs Y. No delays are required, although the availability of delays might ease the implementation.

This opens a perspective of signal processing possibilities with selectivity beyond that of the cochlear filter, an idea exploited by other hybrid models such as the lateral inhibitory network (LIN) of Shamma (1985) or the phase opponency of Carney et al. (2002), or earlier proposals of a “second filter” (Huggins and Licklider, 1951), or the more recent SFIE model of Nelson and Carney (2004), or the “synthetic delay lines” of de Cheveigné and Pressnitzer (2006). The interplay between cochlear filtering and time-domain processing is integral to all these models.

This perspective assumes perfect linearity, whereas we know that transduction and neural processing have imperfect linearity and limited dynamic range. Indeed, measurements of binaural or harmonicity-based unmasking (reviewed in de Cheveigné, 2021) suggest that the benefit of post-cochlear filtering might be limited to about 3-15 dB. Cochlear filtering is required to achieve a greater attenuation, and thus the concepts of “critical band” and “resolvability” remain entirely relevant. These hypotheses warrant further investigation.

#### 5. Variants of the cancellation filter

A partial of frequency f can be cancelled with a delay equal to any multiple T= k/fof its period. There might be an advantage of choosing a larger rather than smaller value of k, as the resulting filter imposes less attenuation to the spectral region immediately adjacent the peak of the peripheral filter, as illustrated in Fig. 15. This requires a larger value of T.

**Fig. 15.**
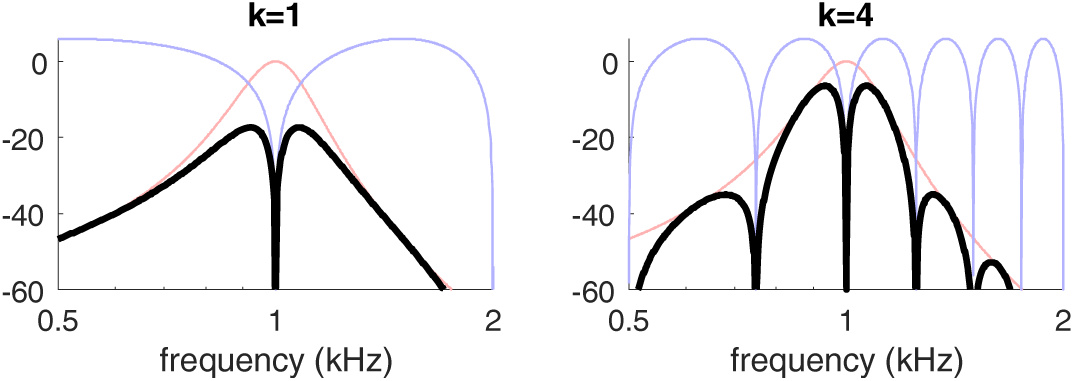
Transfer functions of compound filter with CF=1 kHz for T = 1 ms (left) or T = 4 ms (right). Both offer perfect rejection at 1 kHz, but the filter on the right imposes less attenuation on the spectral region near 1 kHz.

On the other hand, it has been argued that there may be a penalty for longer delays in the auditory system (as reviewed by de Cheveigné and Pressnitzer, 2006), in which case k = 1 is preferable. Delays might be further shortened by noting that many channels pass at most two adjacent partials of a harmonic sound (Fig. 3). Their output can be supressed by applying a cascade of two cancellation filters tuned to those partials. These variants are worth considering when simulating the model or searching for its correlates within the brain.

## V. Conclusion

This paper considered the hypothesis of a simple filtering mechanism implemented at the output of the cochlea for the purpose of ensuring invariance with respect to maskers that are harmonic, locally harmonic, or spectrally sparse. The model accounts qualitatively for the masking asymmetry between pure tone and narrowband noise, and between harmonic and inharmonic complex tones. The filter involves a channel-specific delay parameter that is estimated automatically within each channel. As a spinoff of this unmasking mechanism, the delay parameter might also serve as a cue to pitch, or to detect frequency change between inharmonic stimuli that do not evoke a clear pitch. The model was discussed as an example of a wider class of putative processing mechanisms that associate quasi-linear filtering in the cochlea with time-domain neural processing in the brainstem.

## Acknowledgments

This work was supported by grants ANR-10-LABX-0087 IEC, ANR-10-IDEX-0001-02 PSL, and ANR-17-EURE-0017.

